# Predicting cell types in single cell mass cytometry data

**DOI:** 10.1101/316034

**Authors:** Tamim Abdelaal, Vincent van Unen, Thomas Höllt, Frits Koning, Marcel J.T. Reinders, Ahmed Mahfouz

**Affiliations:** Delft Bioinformatics Lab, Delft University of Technology, 2628 CD Delft, The Netherlands; Leiden Computational Biology Center, Leiden University Medical Center, 2333 ZC Leiden, The Netherlands; Department of Immunohematology and Blood Transfusion, Leiden University Medical Center, 2333 ZA Leiden, The Netherlands; Computer Graphics and Visualization, Delft University of Technology, 2628 CD Delft, The Netherlands

## Abstract

**Motivation:** Mass cytometry (CyTOF) is a valuable technology for high-dimensional analysis at the single cell level. Identification of different cell populations is an important task during the data analysis. Many clustering tools can perform this task, however, they are time consuming, often involve a manual step, and lack reproducibility when new data is included in the analysis. Learning cell types from an annotated set of cells solves these problems. However, currently available mass cytometry classifiers are either complex, dependent on prior knowledge of the cell type markers during the learning process, or can only identify canonical cell types.

**Results:** We propose to use a Linear Discriminant Analysis (LDA) classifier to automatically identify cell populations in CyTOF data. LDA shows comparable results with two state-of-the-art algorithms on four benchmark datasets and also outperforms a non-linear classifier such as the k-nearest neighbour classifier. To illustrate its scalability to large datasets with deeply annotated cell subtypes, we apply LDA to a dataset of ~3.5 million cells representing 57 cell types. LDA has high performance on abundant cell types as well as the majority of rare cell types, and provides accurate estimates of cell type frequencies. Further incorporating a rejection option, based on the estimated posterior probabilities, allows LDA to identify cell types that were not encountered during training. Altogether, reproducible prediction of cell type compositions using LDA opens up possibilities to analyse large cohort studies based on mass cytometry data.

**Availability:** Implementation is available on GitHub (https://github.com/tabdelaal/CyTOF-Linear-Classifier).

## 1 Introduction

Mass Cytometry by time-of-flight (CyTOF) is a valuable tool for the field of immunology, as it allows high-resolution dissection of the immune system composition at the cellular level (Bandura et al., 2009). Advances in CyTOF technology provide the simultaneous measurement of multiple cellular protein markers (> 40), producing complex datasets which consist of millions of cells (Spitzer and Nolan, 2016). Many recent studies have shown the utility of CyTOF to identify either canonical or new cell types while profiling the immune system. These include 1) the characterization of cell type heterogeneity for a specific cancer (Amir et al., 2014; Levine et al., 2015; Chevrier et al., 2017), 2) assigning signature cell populations when profiling a specific disease (van Unen et al., 2016), and 3) monitoring the immune system response to various infections (Newell et al., 2012, 2013).

A key step in mass cytometry analysis is the accurate identification of cell types in a given sample. The high number of dimensions in CyTOF data has forced researchers to depart from manual gating strategies based on two-dimensional plots because it’s very labour intensive and subjective (Newell and Cheng, 2016). These limitations greatly impede the translational aspects of these technologies. Major efforts have been made to facilitate the analysis of CyTOF data by means of clustering (unsupervised learning) methods. These include SPADE (Qiu *et al.*, 2012), Phenograph (Levine *et al.*, 2015) and VorteX (Samusik *et al.*, 2016), and they are often combined with dimensionality reduction methods like PCA (Hotelling, 1933), t-SNE (van der Maaten and Hinton, 2008; Pezzotti *et al.*, 2017), and HSNE (Pezzotti *et al.*, 2016; Van Unen *et al.*, 2017).

While very instrumental in analysing high-dimensional data, clustering approaches require substantial input from researchers to assign cell type labels. This can be done by visually exploring the data, either by gating the biaxial marker expression scatter plots in the case of flow cytometry (FC), or by overlaying the marker expression profiles on a low-dimension representation (e.g. tSNE map) and define the different cell types (clusters). This follows from clustering being an ill-defined problem, which is even more pronounced with increasing number of cells in CyTOF datasets. While useful in identifying ‘new’ cell types in explorative experiments, manual interventions also significantly limit the reproducibility of identifying cell-types across different (batches of) samples. The latter is particularly important in studies comparing different conditions, for example in cohort studies, and when building predictors based on cell compositions, which cannot be performed using clustering methods.

These limitations are inherent to both FC and CyTOF, albeit more pronounced in the later given the higher number of dimensions and the larger number of cells being measured. In the field of FC, several supervised approaches have been proposed to automatically identify cell populations. They have been shown to match the performance of centralized manual gating based on benchmark datasets from challenges organized by the FlowCAP (“Flow Cytometry: Critical Assessment of Population Identification Methods”) Consortium (Hsiao *et al.*, 2016; Lux *et al.*, 2018). These approaches rely on learning the manual gating from a set of training samples, and transferring the learned thresholds for the gates to new test samples.

As gating is done based on two dimensional views of the data, this is not a feasible approach anymore for CyTOF data, since the number of markers is generally around 40, resulting in ~ 2^40^ of gates that need to be defined (at least one for every pair of markers). Moreover, manual gating generally assumes that cells of interest can be selected for by dichotomizing each marker, i.e. splitting cells on the basis of a marker being positively or negatively expressed (identified by a threshold value, i.e. the gate). However, analyses of CyTOF data have repeatedly shown that cell type composition is much more complex, showing many clusters that are described by a combination of all marker expressions (Van Unen *et al.*, 2017), requiring the need for a multitude of gates that increases the complexity of gating even further.

Consequently, for CyTOF data, alternative gating approaches need to be considered. Recently, two methods have been developed: Automated Cell-type Discovery and Classification (ACDC) (Lee *et al.*, 2017) and DeepCyTOF (Li *et al.*, 2017). ACDC integrates prior biological knowledge on markers of specific cell populations, using a cell-type marker table in which each marker takes one of three states (1: positively expressed, −1: negatively expressed, 0: don’t care) for each cell type. This table is then used to guide a semi-supervised random walk classifier of canonical cell types (i.e. cell types with defined marker expression patterns). DeepCyTOF applies deep neural networks to learn the clustering of one sample, and uses the trained network to classify cells from different samples. Both methods achieve accurate results on a variety of datasets. However, both methods rely on sophisticated classifiers. Interestingly, none of these methods compared their performance to less complex classifiers. Further, both methods focused mainly on classifying canonical cell types, which is not the main focus of CyTOF studies which usually relies on the large number of markers measured for deep interrogation of cell populations.

In this work, we show that a linear discriminant analysis (LDA) classifier can accurately classify cell types in mass cytometry datasets. Compared to previous methods, LDA presents a simpler, faster and reliable method to assign labels to cells. Moreover, using LDA instead of more complex classifiers enables the analysis of large datasets comprised of millions of cells. To illustrate this, we tested the applicability of LDA in classifying not only the canonical cell types but also when deeper subtyping the human mucosal immune system across multiple individuals, where the classification task become harder as the differences between cell types is much smaller.

## 2 Methods

We define a *cell* as the single measurement event in CyTOF data, c ∈ ℛ^*P*^ where *p* is the number of markers on the CyTOF panel. Cells are being measured collectively from one *sample*, which is the biological specimen collected from an individual. A sample usually consists of thousands of cells, i.e. 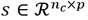, where *n_c_* is the number of cells in sample *s*. A CyTOF *dataset* consists of multiple samples, 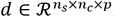, where *n_s_* is the number of samples in the dataset that can comprise different groups of patients. Ultimately, we are interested in identifying a *cell type* which is a collection of cells that have a similar protein marker expression. Usually the different cell types are derived from clustering a large collection of cells collected from different samples using an unsupervised clustering approach. Consequently, we will also refer to cell types as clusters or a cell population.

### 2.1 Datasets description

We used four public benchmark datasets to evaluate our classifier, for which manually gated populations were available and used as ground truth reference (Table 1 summarizes the datasets used in this study). First, the **AML dataset** is a healthy human bone marrow mass cytometry dataset (Levine *et al.*, 2015), consisting of 104,184 cells analysed using 32 markers resulting in 14 cell type populations defined by manual gating. Second, the **BMMC dataset** is also a healthy human bone marrow dataset (Bendall *et al.*, 2011; Levine *et al.*, 2015), consisting of 81,747 cells analysed with 13 markers, and 24 manually gated populations. Third, the **PANORAMA dataset** entails 10 replicates of mice bone marrow cells (Samusik *et al.*, 2016), analysed using a mass cytometry panel of 39 markers and manually gated into 24 classes, with a total number of cells around 0.5 million. Finally, the **Multi-Center study dataset** is a collection of 16 samples drawn from a single subject (Li *et al.*, 2017), where the first eight samples are collected at the same time and analysed with the same instrument, and the last eight samples are collected two months later and analysed with a different instrument. It contains ~930,000 cells, analysed with 26 markers, where only eight markers were used for the manual gating process (Li *et al.*, 2017), resulting in four canonical cell populations in addition to a fifth class representing the unlabelled cells. Measured expressions were transformed using hyperbolic arcsin with a cofactor of 5 before any further processing for all datasets.

**Table 1.** Summary of the datasets used in this study.

| Dataset | Samples | Cells | Markers | Cell types |
| --- | --- | --- | --- | --- |
| AML | n.a. | 104,184 | 32 | 14 |
| BMMC | n.a. | 81,747 | 12 | 24 |
| PANORAMA | 10 | 514,386 | 39 | 24 |
| Multi-Center | 16 | 929,685 | 8 | 5 |
| HMIS-1 | 47 | 3,553,596 | 28 | 6 |
| HMIS-2 | 47 | 3,553,596 | 28 | 57 |

n.a. = not available

In addition to the benchmark datasets, we used data that we collected from patients with gastrointestinal diseases as well as controls. This Human Mucosal Immune System mass cytometry (HMIS) dataset (van Unen *et al.*, 2016) consists of 102 samples: 47 Peripheral Blood Mononuclear Cells (PBMC) and 55 gut tissue samples. We focused on the PBMC samples only, which are further divided into 14 control samples, 14 samples with Crohn’s Disease (CD), 13 samples with Celiac Disease (CeD) and 6 samples with Refractory Celiac Disease Type II (RCDII). There are ~3.5 million cells in the 47 samples, which are measured with a panel of 28 markers. To annotate cells with cell type information, we clustered all cells across all samples simultaneously using Cytosplore^+HSNE^ (www.cytosplore.org) (Höllt *et al.*, 2016). Cytosplore^+HSNE^ is specifically designed to interactively explore millions of cells. For that it makes use of different (hierarchical) abstraction layers of the individual cells. The bottom layer represents individual cells. To abstract those to the next layer, landmark cells are selected that are cells with a dense neighbourhood. This abstraction is iteratively repeated, with every new layer containing less and less landmark cells, until a top layer that then gives a summarized overview of all cells. At any layer, (landmark) cells are embedded into a two dimensional map using t-SNE (van der Maaten and Hinton, 2008; Pezzotti *et al.*, 2017), and clustered using a density-based Gaussian Mean Shift (GMS) clustering algorithm (Comaniciu and Meer, 2002).

For the HMIS dataset we constructed three layers. For the top (overview) layer, we annotated the clusters into six major immune lineages on the basis of the expression of known lineage marker: 1) CD4+ T cells, 2) CD8+ T cells, including TCRgd cells, 3) B cells, 4) CD3-CD7+ Innate Lymphocytes (ILCs), 5) Myeloid cells, and 6) Others, representing unknown cell types (Fig.1). Note that although the clusters are created from the abstracted landmark cells, the cluster annotations are propagated to every individual cell, resulting in ~3.5 million cells annotated into six canonical cell types, which we denoted the **HMIS-1 dataset**. Next, in order to find subtypes at a more detailed level, we explored one layer down for each of the six cell types separately, producing six separate t-SNE maps (Fig. 1). For each map, we applied GMS clustering with a kernel size of 30 (default value). For each cluster, we calculated a cluster representation by taking the median expression of each marker for all individual cells annotated with that cluster. We automatically merged clusters when the correlation between cluster representatives is above 0.95. We discarded clusters containing less than 0.1% of the total number of cells (< 3500 cells). In total we ended up with 57 cell types (11 CD4+ T cells, 9 CD8+ T cells, 4 TCRgd cells, 11 B cells, 11 CD3-CD7+ ILCs, 6 Myeloid cells and 5 Others) for the ~3.5 million cells, which we denoted the **HMIS-2 dataset**. Cell counts per cell type and per sample are summarized in Supplementary Fig. S1.

**Fig. 1.**
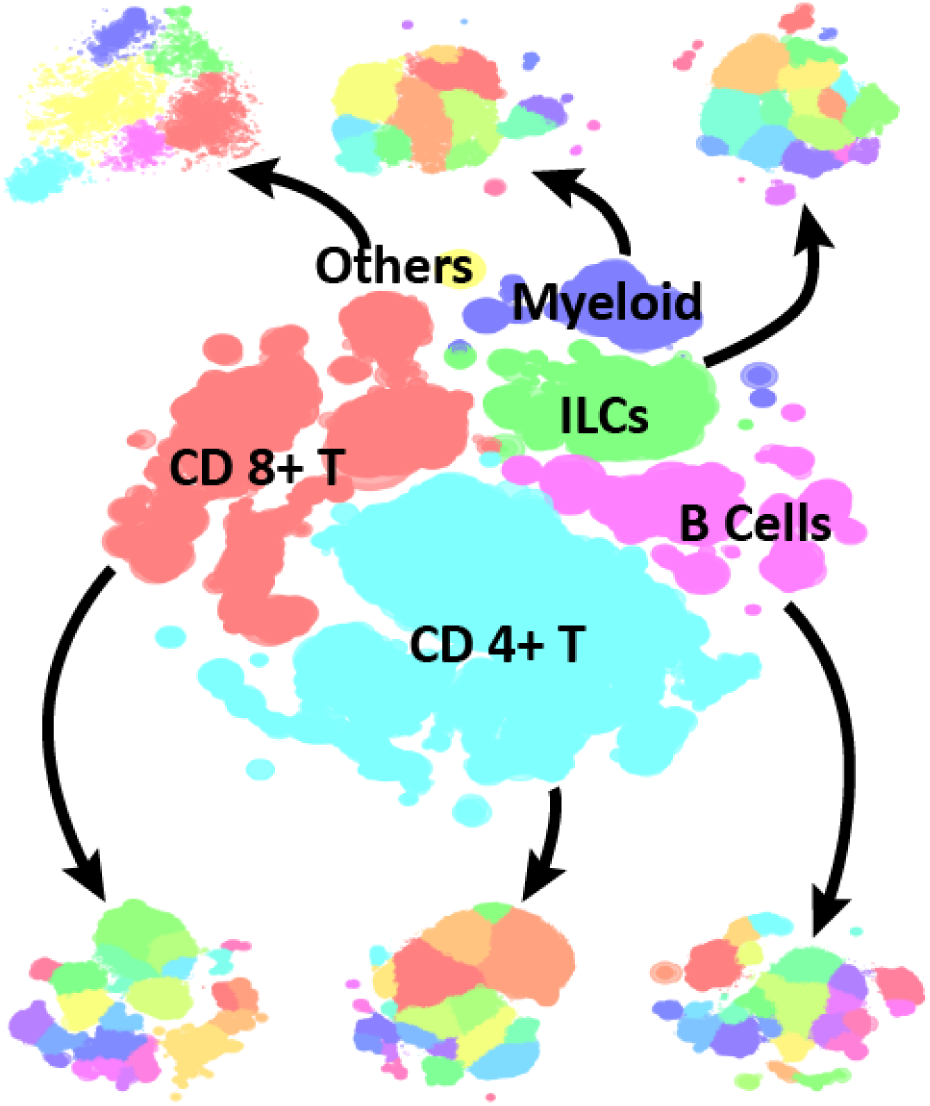
Annotation of cells in the HMIS dataset. The middle image shows the embedding of the overview (top) HSNE layer, clustered into six major cell types. Next, a separate tSNE map is obtained per cell type by exploring one layer down.

### 2.2 Cell type predictor

To determine cell types in a newly measured sample, one would need to re-cluster the new sample with all previous samples. Besides being a tedious task, cells from the new sample will influence the clustering and by that change the previously identified cell types, severely hampering reproducibility. Therefore, we learn the different cell types from a training set with annotated cells. The cell types in the new sample can then simply be predicted by this learned cell-type predictor.

We propose to use a (simple) Linear Discriminant Analysis (LDA) classifier (fitcdiscr function Matlab R2016b) as well as a k-NN classifier to check whether the performance of a non-linear classifier would outperform the linear LDA classifier. For the k-NN classifier, we used the Matlab R2016b implementation (fitcknn function, with Euclidean distance and k = 50 neighbours). We adopted an editing approach to train the k-NN classifier to reduce the training set size (and consequently keep computation times reasonable). The editing is done according to the following pseudo code.

~~~
*Temp_Training* ⟵ random 50,000 cells from *Training_Data*
**while** (not all *Training_Data* is covered)
    *Temp_Testing* ⟵ another random 50,000 cells from *Training_Data*
    Apply prediction on *Temp_Testing* and add misclassified cells to *Temp_Training*
    *Temp_Training* ⟵ *Temp_Training* + Misclassified from *Temp_Testing*
**end while**
*Final_Training* ⟵ *Temp_Training*
~~~

### 2.3 Performance metrics

To evaluate the quality of the classification, we used four metrics:

1. The classification accuracy (fraction of correctly identified cell).
2. The F1-score (harmonic mean of the precision and recall) for which we report the median value across all cell types. When comparing to DeepCyTOF (Li *et al.*, 2017), we use the weighted average of F1-scores per cell type size, to produce a fair comparison.
3. The maximum difference in population frequencies, defined as 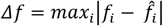where *f_i_* and *f̂_i_* represents the true and the predicted percentage cell frequencies for the *i*-th cell population, respectively.
4. The Root of Sum Squared Error (RSSE) per sample and per cell type, defined as 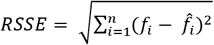. In case of measuring the error per sample, *f_i_* and *f̂_i_* represents the true and the predicted percentage cell frequencies, respectively, for the *i*-th cell population per sample, and *n* = *n_t_* (total number of cell types). In case of measuring the error per cell type, *f_i_* and *f̂_i_* represents the true and the predicted percentage cell frequencies, respectively, for a certain cell type in the *i*-th sample, and *n* = *n_s_* (total number of samples).

### 2.4 Performance estimation

The performance of a classifier is evaluated using three different cross-validation setups:

1. *CV-Cells*: 5-fold cross validation applied over all the cells.
2. *CV-Samples*: A leave-sample-out cross validation over all the samples, regardless of the number of cells within each sample. The classifier is trained using the cells of the samples in the training set, then the cell type prediction is done per left-out sample.
3. *Conservative CV-Samples*: Similar to *CV-Samples*, but with the main difference that the ground-truth reference labels, acquired by clustering, are not used for training. Instead, for each set of training samples the data is re-clustered, resulting in new cell types. These new cell types are then used to train the classifier, which is subsequently used to predict the labels of the cells of the left-out sample. Since the labels of the training set and the ground-truth are now different, we matched the cluster labels by calculating their pairwise correlation (Pearson’s *r*) using the median marker expression of each cluster. Each training cluster is matched to the ground-truth cluster with which the correlation is maximum.

For the AML and the BMMC datasets, we evaluated the performance using the *CV-Cells* setup only, since no sample information is provided. For the PANORAMA and Multi-Center datasets, we used both the *CV-Cells* and *CV-Samples* setups, since we have the sample information. Considering the number of samples in each dataset, we used a 5-fold *CV-Samples* for the PANORAMA dataset and a 4-fold *CV-Samples* for the Multi-Center dataset. For the HMIS-1 and HMIS-2 datasets, we used all three cross validation setups, using a 3-fold *CV-Samples* and *Conservative CV-Samples*.

### 2.5 Rejection Option

To be able to detect new cell types, we decided to include a rejection option for LDA by defining a minimum threshold for the posterior probability of the assigned cell types. Thus, a cell is labelled as ‘unknown’ whenever the posterior probability is less than a predefined threshold set.

### 2.6 Feature Selection

To avoid overfitting, we explored the need to reduce the number of markers (i.e. features) by applying feature selection on the training data. First, we applied a 5-fold *CV-Cells* and used the classification performance for every individual marker on the training data to rank all markers in a descending order. Next, we applied another 5-fold *CV-Cells* on the training data and trained as many classifiers as there are markers. The first classifier is based on the top marker only, the second one on the two top ranked markers, etc. Then we select the classifier which generates the best cross validation performance over the training set. This classifier is subsequently tested on the test set and the performance is reported.

## 3 Results

### 3.1 LDA has comparable performance on CyTOF data to complex classification approaches

To evaluate the performance of the LDA classifier, we compared LDA with two recent state-of-the-art methods for classifying CyTOF data, ACDC (Lee *et al.*, 2017) and DeepCyTOF (Li *et al.*, 2017). We used the AML, BMMC and PANORAMA datasets (used by ACDC) and the Multi-Center dataset (the only available dataset used by DeepCyTOF). We compared the performance of LDA with the reported values in these two studies (Table 2).

**Table 2.** Performance summary of LDA versus ACDC and DeepCyTOF

|  | LDA<br>*CV-Cells* | LDA<br>*CV-Samples* | ACDC | Deep-CyTOF |
| --- | --- | --- | --- | --- |
| Accuracy |  |  |  |  |
| AML | 98.13 ± 0.09 | n.a. | 98.30 ± 0.04 | n.a. |
| BMMC | 95.82 ± 0.10<br>(95.61 ± 0.16)* | n.a. | 92.90 ± 0.5 | n.a. |
| PANOR-AMA | 97.16 ± 0.07<br>(97.70 ± 0.03)* | 97.22 ± 0.31<br>(97.67 ± 0.29)* | n.r. | n.a. |
| Multi-Center | 96.98 ± 0.03 | 96.91 ± 1.17<br>(96.90 ± 3.26)** | n.a. | n.r. |
| Median F1-score |  |  |  |  |
| AML | 0.95 | n.a. | 0.93 | n.a. |
| BMMC | 0.85<br>(0.85)* | n.a. | 0.60 | n.a. |
| PANOR-AMA | 0.93<br>(0.95)* | 0.93<br>(0.95)* | 0.88 | n.a. |
| Multi-Center*** | 0.95 | 0.95<br>(0.93)** | n.a. | 0.93 |

n.a. = not available, n.r. = not reported.
\*Four classes considered unknown, similar to ACDC.
\*\*Only one sample is training (Sample 2), similar to DeepCyTOF.
\*\*\*Weighted F1-score.

Since there was no sample information available for the AML and BMMC datasets, we evaluated the performance of the LDA classifier on both datasets using the *CV-Cells* setup only. For the AML dataset, LDA achieved comparable performance in terms of accuracy and median F1-score to ACDC. For the BMMC dataset, we applied the LDA classifier to classify all 24 cell populations, resulting in ~ 96% accuracy and 0.85 median F1-score. To have a fair comparison with ACDC, we also considered four classes as unknown (Lee *et al.*, 2017) then classified only 20 cell classes. In both cases, LDA outperformed ACDC, specially based on the median F1-score.

On the PANORAMA dataset, we tested the LDA classifier to classify all 24 populations using both the *CV-Cells* and *CV-Samples* setups. In addition, we tested the performance of LDA on 22 populations only to have a fair comparison with ACDC (Lee *et al.*, 2017). In both cases LDA produces relatively high accuracy and median F1-score, and repeatedly outperformed ACDC in terms of the median F1-score (no accuracy reported by ACDC).

For the Multi-Center dataset, we applied *CV-Cells* and *CV-Samples* yielding an accuracy of ~97% and weighted F1-score of 0.95 for both setups. To have a fair comparison with DeepCyTOF, we only used sample no. 2 for training and tested the classifier performance on the other 15 samples. LDA shows a similar performance profile per sample as DeepCyTOF, where the weighted F1-score is relatively high for samples 1-8 (samples measured on the same instrument as sample no. 2), and lower performance for samples 9-16 (Supplementary Fig. S2). Overall, LDA had a comparable weighted F1-score to the DeepCyTOF, excluding the calibration step (Li *et al.*, 2017).

### 3.2 LDA accurately classifies immune cells in a larger dataset with deeper annotation of cell subtypes

To test our hypothesis that LDA can achieve acceptable performance on large datasets and with more detailed cell subtyping, we applied LDA to the HMIS dataset comprised of ~3.5 million cells. The HMIS data was clustered at two levels of detail (see Methods) resulting in two different annotations for the HMIS data set: HMIS-1, representing six major lineages, and HMIS-2 containing 57 cell types. For both annotations, we applied all three cross validation setups, *CV-Cells*, *CV-Samples* and *Conservative CV-Samples* (Table 3).

**Table 3.** Performance summary of LDA on the HMIS dataset.

|  | LDA<br>*CV-Cells* | LDA<br>*CV-Samples* | LDA *Conservative*<br>*CV-Samples* |
| --- | --- | --- | --- |
| Accuracy |  |  |  |
| HMIS-1 | 99.38 ± 0.01 | 99.02 ± 2.26 | 98.91 ± 1.87 |
| HMIS-2 | 87.19 ± 0.05 | 86.11 ± 3.86 | 78.69 ± 8.65 |
| Median F1-score |  |  |  |
| HMIS-1 | 0.99 | 0.99 | 0.99 |
| HMIS-2 | 0.81 | 0.79 | 0.62 |

We first tested the LDA performance on HMIS-1, hence only classifying the canonical cell types. LDA achieved an accuracy > 99% and a median F1-score > 0.98 for both *CV-Cells* and *CV-Samples*. Next, we applied LDA to HMIS-2, which implied classifying cells into 57 different cell types including abundant and rare cell populations. As expected, LDA had a lower performance on HMIS-2 compared to HMIS-1 using both *CV-Cells* and *CV-Samples*. The confusion matrix in Fig. 2A shows that the performance drop between HMIS-1 and HMIS-2 is mainly caused by misclassifications within the same major lineages. We further investigated the LDA performance across different sample types (Control, CeD, RCDII and CD) in the HMIS dataset. Fig. 3A shows that LDA has the highest accuracy for the control samples, while the lowest accuracy is for the RCDII samples.

**Fig. 2.**
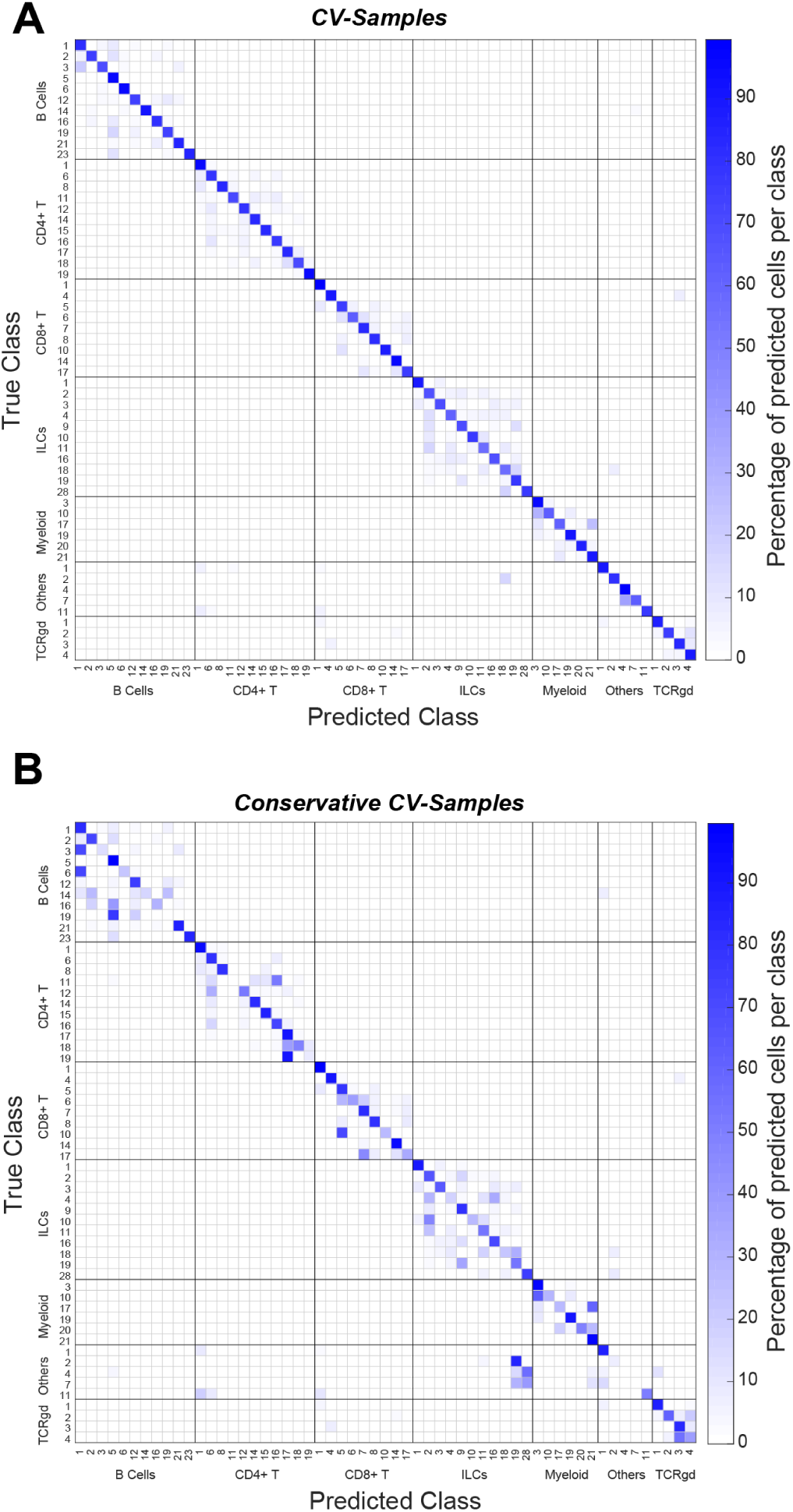
LDA performance on the HMIS-2 dataset. **(A)**Classification confusion matrix when using *CV-Samples* setup, showing high percentages along the matrix diagonal, as well as that most of the misclassification (off-diagonal values) falls within the major cell types. **(B)** Classification confusion matrix when using *Conservative CV-Samples* setup, showing lower percentages along the matrix diagonal compared to the *CV-Samples* setup. Each cell (square) in the confusion matrix represents the percentage of overlapping cells between true and predicted class.

**Fig. 3.**
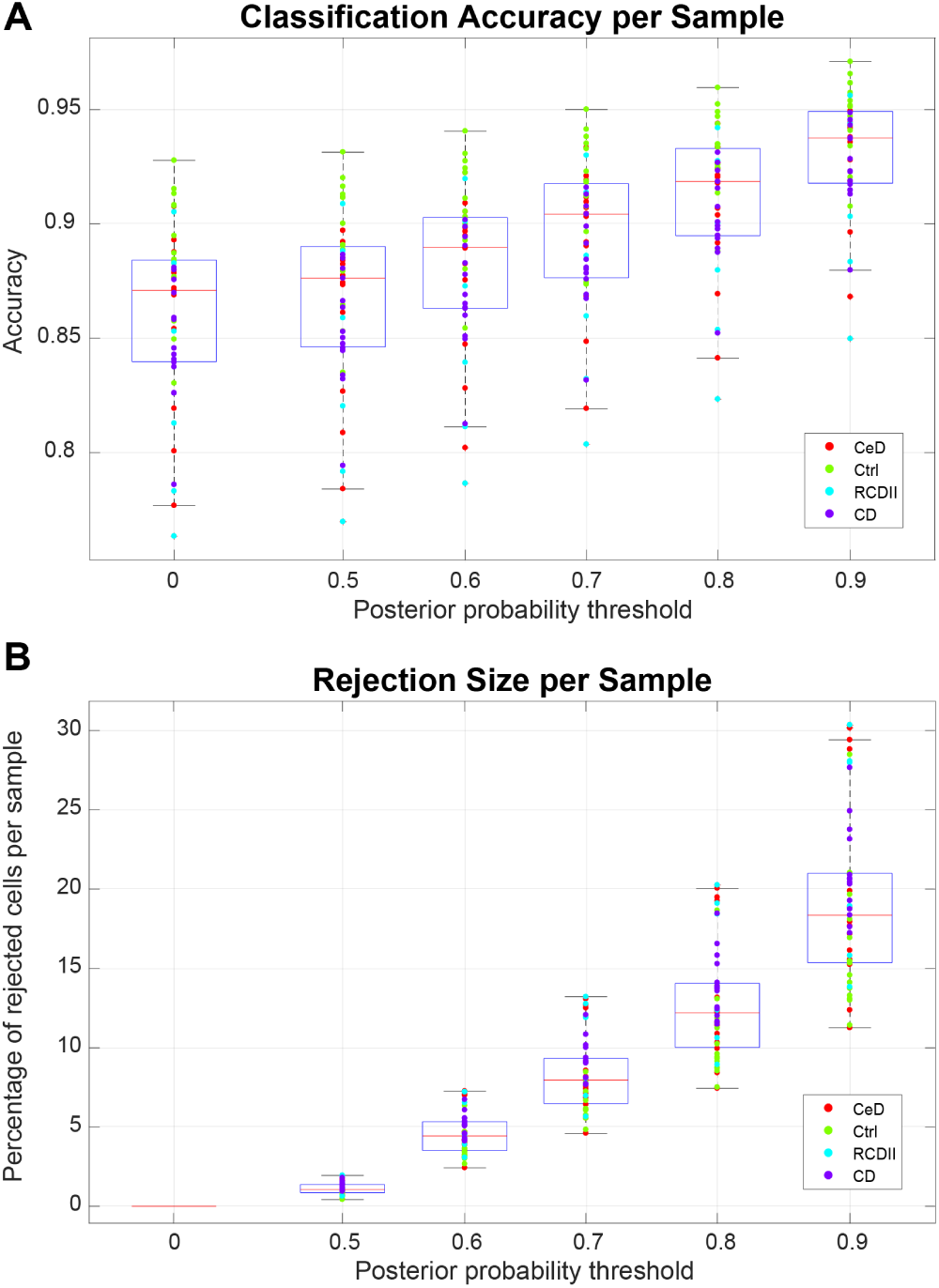
LDA accuracy and rejection size per sample. **(A)** boxplot of the LDA accuracy distribution per sample, while using a rejection threshold, (0 = no rejection). **(B)** boxplot of the rejection percentage per sample while using a rejection threshold, (0 means no rejection). Each dot represents a sample coloured according to the sample type (CeD = Celiac Disease, Ctrl = Control, RCDII = Refractory Celiac Disease Type II, CD = Crohn’s Disease).

To better mimic a realistic scenario and avoid any leakage of information from the testing samples by considering all samples when preclustering cells to determine the ground truth labels, we used a *conservative CV-Samples* setup to evaluate the LDA classifier (see Methods). For the HMIS-1 dataset representing the major lineages, the performance of LDA in the *Conservative CV-Samples* was comparable to the other setups (*CV-Cells* and *CV-Samples*), Table 3. The performance of the LDA classifier dropped when considering the *Conservative CV-Samples* setup on HMIS-2 that contains a multitude of cell types. However, the lower performance can be explained by miss-matching clusters between the training set and the ground-truth, which introduces classification errors. For example, cluster ‘CD4 T 11’ is never predicted by the classifier, which means all cells falling within this cluster will be misclassified (Fig. 2B). This is because in all 3-folds, no training cluster matches to this ground-truth cluster ‘CD4 T 11’ (Supplementary Fig. S3). Whereas in case of HMIS-1, with only six dissimilar clusters, the clusters map works perfectly, resulting in high performance (Supplementary Fig. S4).

### 3.3 LDA accurately estimates cell population frequencies

One of the main aims of CyTOF studies is to estimate the frequencies of different cell types in a given sample. We evaluated the LDA prediction performance in terms of predicted population frequencies, by calculating the maximum difference in population frequencies, ∆*f*, for each dataset (see Methods). LDA produced comparable population frequencies to the manually gated populations (Fig. 4). The maximum difference in population frequency (∆*f*) was 0.4%, 0.65%, 0.64% and 1.1% for the AML, BMMC, PANORAMA and the Multi-Center datasets, respectively.

**Fig. 4.**
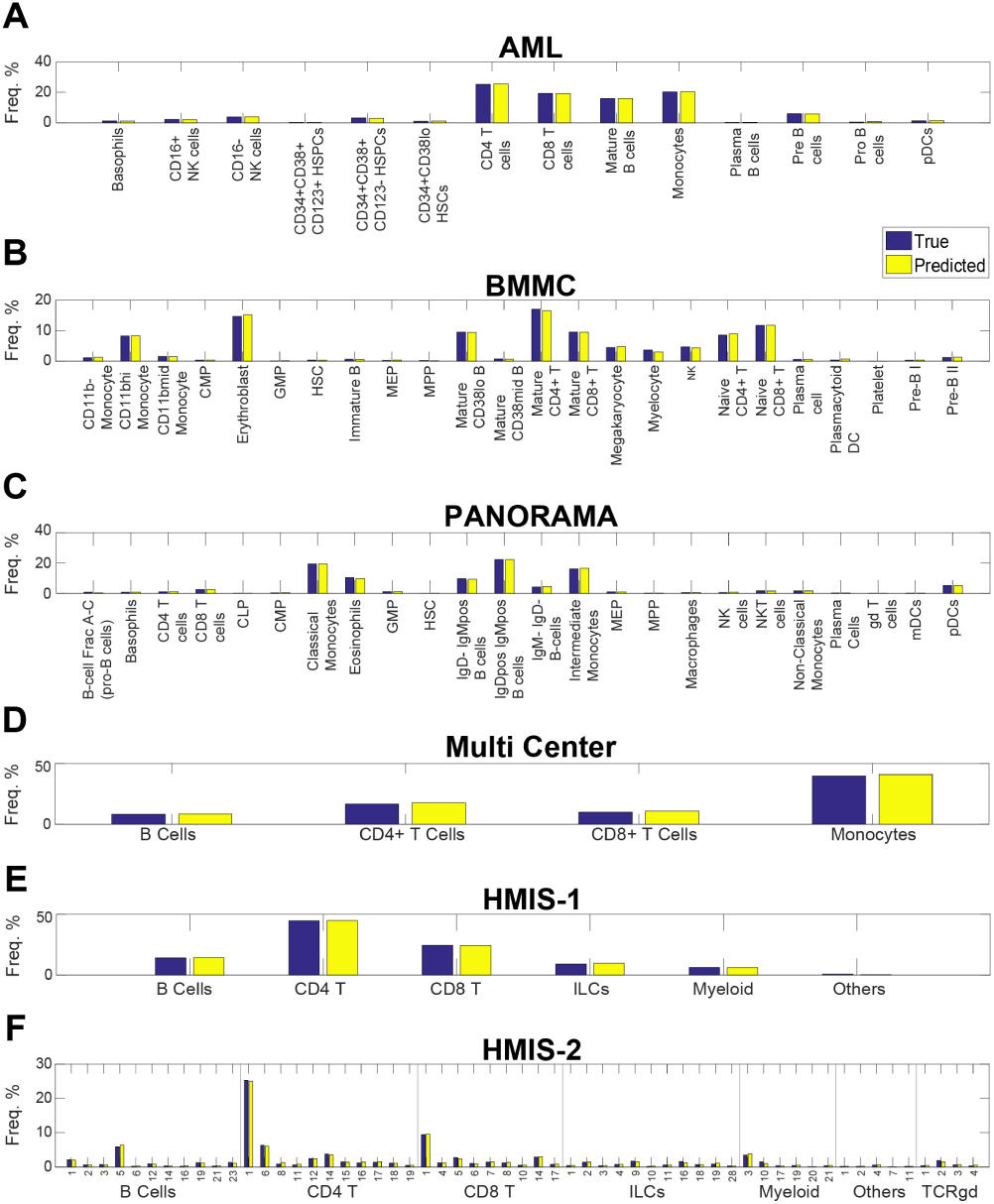
True vs predicted population frequencies. **(A)** AML, **(B)** BMMC, **(C)** PANORAMA, **(D)** Multi-Center, **(E)** HMIS-1 and **(F)** HMIS-2.

For the HMIS-1 dataset, LDA has ∆*f* of 0.59% across the six major cell types. Interestingly, despite the drop in the accuracy of predicting cell labels on HMIS-2 compared to HMIS-1, the population frequencies are not significantly affected. The maximum difference of population frequencies in HMIS-2 was 0.46% among all 57 cell types (Fig. 4F). This small ∆*f* shows that LDA produces accurate performance with respect to the ground-truth reference, even at a detailed annotation level.

We investigated the population differences per sample and per cell type using the *CV-Samples* setup in the HMIS-2 dataset, by calculating the average squared differences between the estimated and true frequencies (RSSE, see Methods). We obtained small RSSE values with a maximum of 0.074 (sample no. 10) and 0.082 (‘Myeloid 10’ cluster) across different samples and different cell types, respectively (Supplementary Fig. S5). For sample no. 10, the maximum absolute population difference was 5.17% for ‘Myeloid 3’ cell type. For ‘Myeloid 10’ cluster, the maximum absolute difference 5.12% across all cells.

### 3.4 LDA performs on highly abundant as well as race cell types

To evaluate the performance of LDA for abundant and rare cell populations, we investigated the F1-score per cell type versus the population size. Fig. 5 shows the F1-score for all 57 cell types in the HMIS-2 dataset obtained using the *CV-Samples*. Remarkably, LDA performs well for large cell populations, as well as the majority of the small cell populations, with a median F1-score of 0.7915 for populations that contain less than 0.5% of the total cells.

**Fig. 5.**
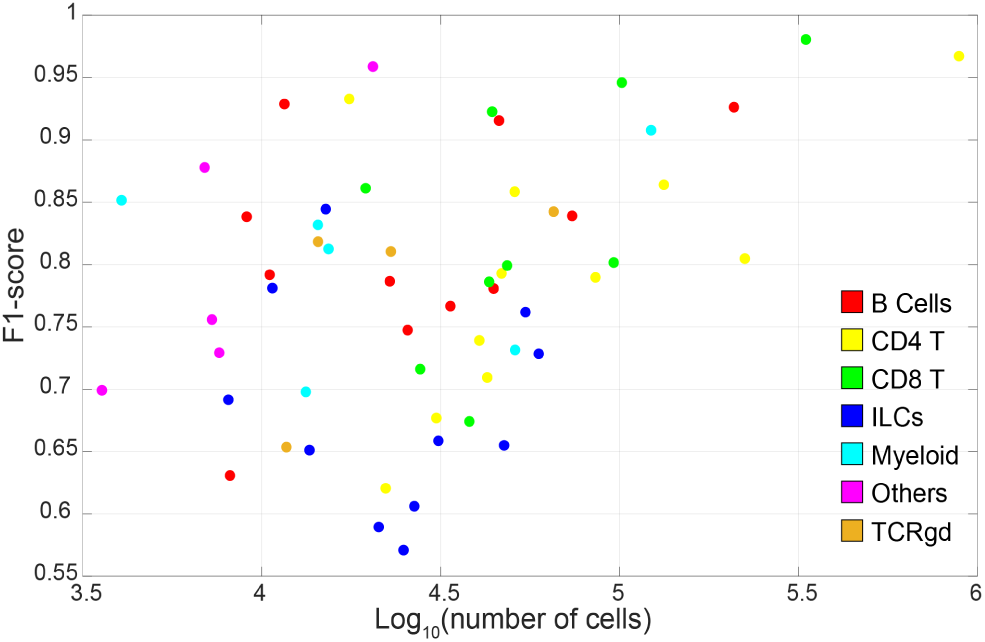
Relationship between performance and population size. Scatter plot of the F1-score vs. the population size (number of cells) for the HMIS-2 dataset evaluated using *CV-Samples*. Each dot represents one cell type and coloured according to the major cell type annotation.

For the *Conservative CV-Samples* setup, the LDA performance is still high for large cell populations, but the F1-score drops for small populations reinforcing that the drop in performance of the *Conservative CV-Samples* is driven by the limitations with the cluster matching rather than the performance of the LDA (Supplementary Fig. S6A). For populations that contain less than 0.5% of the total cells, the median F1-score is 0.4753. Similar patterns were observed for the other four datasets (Supplementary Fig. S6B-E).

### 3.5 LDA as a probabilistic classifier directly allows the detection of unseen cell types

A major advantage of clustering and visual analytics over classification approaches is the ability to identify novel unknown cell types. Here, we show that LDA as a probabilistic classifier can be used to flag unknown cells that do not match any of the training cell types. We incorporated a rejection option to allow the classification of a cell as ‘unknown’ when the posterior probability of the classification of any cell is low. Fig. 3A shows the classification accuracy across samples from the HMIS-2 dataset, after excluding unknown cells for which the posterior probability is lower than a certain threshold. As expected, setting a threshold on the posterior probability resulted in more accurate predictions. For example, setting a threshold at 0.7 resulted in an accuracy of 89.54 ± 3.25% (compared to 86.11 ± 3.86% without any thresholds), while assigning ~8% of cells per sample as unknown.

Further, we observed a reverse pattern between the accuracy of cell classification and the percentage of cells classified as unknown per sample (Fig. 3A and 3B). For instance, LDA has the highest accuracy on classifying cells from the control samples and hence control samples are less likely to entail rejected (unknown) cells. On the other hand, the accuracy is the lowest on RCDII samples which also have the highest rejection percentages. Fig. 3 further shows that both the accuracy and the rejection size increase with increasing the minimum threshold of the posterior probability.

### 3.6 Rejection option targets rare sample-specific cell types

Next, we investigated the effect of the rejection option on rare and abundant cell populations. In the HMIS-2 dataset, the population frequencies of the 57 cell types varied from 25.2% to 0.1% of the total number of cells in the HMIS-2 dataset (Fig. 6A). Further, we observed a variable distribution of cell types across different sample types (control, CeD, RCDII and CD), Fig. 6B. Although the majority of cell types were evenly distributed over all samples, some were disease-specific, especially the rare cell types. Using a rejection threshold of 0.7, we calculated the rejection ratio per cell type per sample (Fig. 6D) as the number of cells assigned as ‘unknown’ of one cell type in one sample, divided by the total number of cells of that cell type in all samples. We compared these rejection ratios with the cell type frequencies over the samples (Fig. 6C) where a value close to 100% means that the cell type is specific to only one sample. We observed a strong correlation between the cell type rejection ratios and the frequencies over the samples (Fig. 6E). For example, the majority of ‘Others 2’ (83.87%) comes from one CeD sample, within which ‘Others 2’ is prominently present (7.44% of the cells in this sample belong to ‘Others 2’ Supplementary Fig. S1). The classifier rejects ~15% of these cells, representing a ~12% rejection ratio of the total number of ‘Others 2’ cells. This is a relatively high rejection percentage compared to other cell types (Fig. 6E).The main reason why there is a large rejection ratio for these cells, is because these cells are mainly present in one sample. When this sample is left out in the *CV-Samples* procedure, during testing these cells are rejected because they are missing in the training data. These results support the validity of using the rejection option to label unknown cells, which are likely to be rare sample-specific populations.

**Fig. 6.**
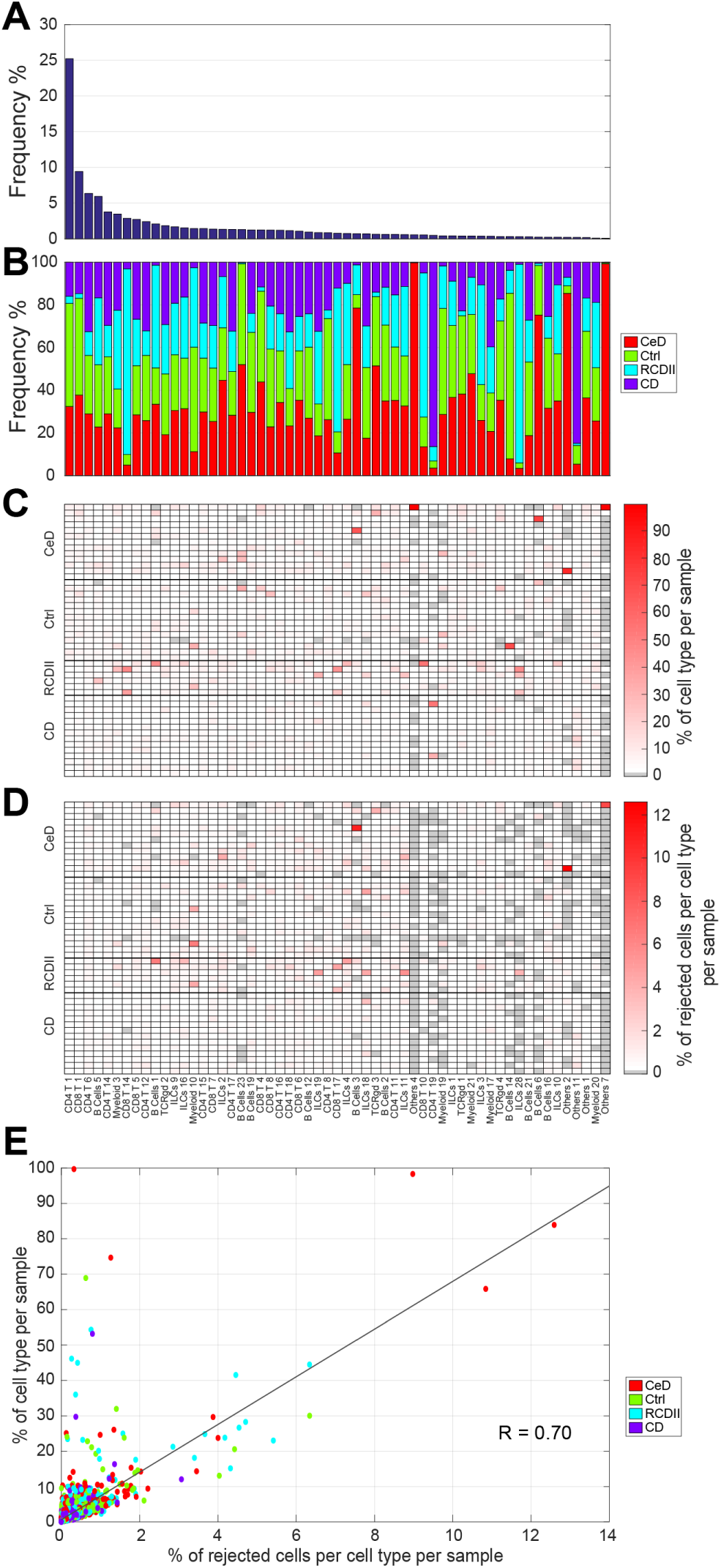
Rejection option effect on variable sized cell populations. **(A)** Cell type frequency across the HMIS-2 dataset, in a descend order. **(B)** Cell type composition in terms of the different sample types (CeD, Ctrl, RCDII, CD). **(C)** Cell type frequencies across samples, normalized by the cell type size across all samples, every column summation is 100%. **(D)** Percentage of rejected cells per cell type per sample, normalized by the cell type size across all samples, using a posterior probability threshold of 0.7. Cell types follow the same order for (A-D). **(E)** Scatter plot between values in (C) and (D) showing a strong correlation of 0.70 between the rejection ratio and the cell type size, per sample. Each point represents a cell type in a particular sample, and points are coloured according to the disease status of the sample annotation.

### 3.7 Linear classification is sufficient for accurate classification of CyTOF data

We have shown that a simple linear classifier such as LDA has a comparable performance to complex non-linear classifiers such as ACDC and DeepCyTOF. To further illustrate that non-linear classification does not perform better than linear classification, we compared the performance of LDA to a k-NN classifier on the HMIS-2 dataset. Again, we found that LDA has a comparable performance to a k-NN classifier with k = 50 (Table 4), suggesting that adding non-linearity to the classification process does not improve performance.

**Table 4.** Performance comparison between LDA and KNN on the HMIS-2 dataset.

|  | Accuracy | Median F1-score |
| --- | --- | --- |
| LDA *Samples CV* | $86.11 \pm 3.86$ | 0.79 |
| k-NN *Samples CV* | $87.73 \pm 4.09$ | 0.81 |
| k-NN *Samples CV FS* | $86.33 \pm 3.17$ | 0.79 |

To reduce the computation time for the k-NN classifier, we employed an editing scheme to reduce the size of the training data (see Methods). Using the proposed editing scheme, we reduced the training data size to an average of 300,000 per training fold (~12 % of the original training set), resulting in a significant speedup of the training and testing times. However, the k-NN classifier still takes on average 180x the time needed by LDA to make predictions for one sample.

Next, we investigated whether feature selection (using less markers during classification) would affect the performance of the classifiers. The k-NN classifier selected only 20 (out of the 28) markers and retained a comparable performance to that obtained using all 28 markers. On the other hand, feature selection did not reduce the number of markers selected by LDA, indicating that LDA requires all the measured markers in order to achieve maximum performance.

## 4 Discussion

In this work, we showed that a linear classifier can be used to automatically assign labels to single cells in mass cytometry data. Using four different CyTOF datasets, we compared the performance of a linear discriminant analysis classifier (LDA) to two recent approaches methods: ACDC (Lee *et al.*, 2017) and DeepCyTOF (Li *et al.*, 2017). Interestingly, LDA achieved similar or better performance compared to ACDC and DeepCyTOF in all four datasets. Compared to ACDC, LDA does not require any additional biological knowledge or assumptions regarding the distribution patterns of markers. Additionally, ACDC requires a cell-type marker table which has several limitations: (1) designing the table can be very challenging in the presence of many cell-types, (2) it is not possible to specify the marker patterns for some cell types (e.g. ACDC ignored 4 subtypes in the BMMC dataset because the table could not be constructed), and (3) the table requires imposing assumptions on the marker distribution (currently binary) which can be challenging to model. Furthermore, results on the BMMC dataset show that LDA can detect rare cell populations having frequencies < 0.5% of the total number of cells, like MPP, HSC, MEP and GMP, which were the main cause of the lower performance of ACDC (Lee *et al.*, 2017). Compared to DeepCyTOF, LDA is a much simpler classifier which means it has substantial advantages with respect to the reduced training/testing times, reproducibility, and scalability to larger datasets.

We further evaluated LDA on a large CyTOF dataset with deep annotation of cell types. We showed that LDA can accurately identify cell types in a challenging dataset of 3.5 million cells comprised of 57 cell types. Further, we showed that the errors made by LDA in assigning cell type labels to each cell has negligible influence on the estimates of cell population frequencies across different individuals.

To show that a linear classifier is indeed sufficient to classify cells in mass cytometry data, we compared LDA to a non-linear classifier (k-NN). We show that the k-NN classifier does not outperform LDA on the HMIS dataset, indicating that there is no added value in using non-linear relationships between the markers. However, when we ran both classifiers with feature selection, LDA required the full set of markers to achieve the best performance. On the other hand, the k-NN classifier was able to achieve the same performance as LDA but using less markers (20 instead of 28). This result suggests that a non-linear classifier might be beneficial to reduce the number of required markers and free valuable slots on the CyTOF panel for additional markers.

For the HMIS dataset, we relied on an initial clustering step to assign ground-truth labels. To avoid any possible leakage of information from the test set of cells by including them into the clustering, we designed a conservative learning scheme. In the conservative scheme, we don’t use the labels obtained by clustering the entire dataset (i.e. ground-truth) for training, but rather re-cluster the training data inside each fold. In addition, this scheme better resembles a realistic scenario in which the new unseen data is never included in the initial assignment of class labels for training. The performance of LDA in this conservative experiment is lower than the initial performance obtained by classical cross validation. However, the drop in performance does not stem from the lack of generalization, as the results show high performance on the overview-level, but rather from the difficulty in matching cluster labels between the ground truth and the training set.

Clustering approaches in general have an advantage over classification methods in that they can be employed to discover new cell types. However, an additional advantage of using a probabilistic classifier such as LDA is that we can directly gain information regarding the accuracy of each decision made by inspecting the posterior probability. We showed that we can allow for a rejection option when the posterior probability of the classification of a particular cell is low. This rejection option can be used to identify ‘unknown’ cells which might require additional investigation to determine their biological relevance. Additionally, we showed that these ‘unknown’ cells are likely to be rare and sample-specific. There is however a trade-off between how confident we are on the correctness of the predictions and the size of the ‘unknown’ class. A stringent threshold (i.e. high posterior probability) means that many cells will be classified as ‘unknown’ which will further require manual investigation.

Taken together, we demonstrated the feasibility of using a simple linear classifier to automatically label cells in mass cytometry data which is a promising step forward to use mass cytometry data in cohort studies.

